# Novel Phosphorylation States of the Yeast Spindle Pole Body

**DOI:** 10.1101/245340

**Authors:** Kimberly K. Fong, Alex Zelter, Beth Graczyk, Jill M. Hoyt, Michael Riffle, Richard Johnson, Michael J. MacCoss, Trisha N. Davis

## Abstract

Phosphorylation regulates yeast spindle pole body (SPB) duplication and separation and likely regulates microtubule nucleation. We report a phosphoproteomic analysis using tandem mass spectrometry of purified *Saccharomyces cerevisiae* SPBs for two cell cycle arrests, G1/S and the mitotic checkpoint, expanding on previously reported phosphoproteomic data sets. We present a novel phosphoproteomic state of SPBs arrested in G1/S by a *cdc4-1* temperature sensitive mutation, with particular interest in phosphorylation events on the γ-tubulin small complex (γ-TuSC). The *cdc4-1* arrest is the earliest arrest at which microtubule nucleation has occurred at the newly duplicated SPB. Several novel phosphorylation sites were identified in G1/S and during mitosis on the microtubule nucleating γ-TuSC. These sites were analyzed *in vivo* by fluorescence microscopy and were shown to be required for proper regulation of spindle length. Additionally, *in vivo* analysis of two mitotic sites in Spc97 found that phosphorylation of at least one of these sites is required for progression through the cell cycle. This phosphoproteomic data set not only broadens the scope of the phosphoproteome of SPBs, it also identifies several γ-TuSC phosphorylation sites influencing microtubule regulation.

## INTRODUCTION

The centrosome is the microtubule organizing center of the cell, responsible for nucleating microtubules and establishing a bipolar spindle during mitosis. In budding yeast, the spindle pole body (SPB) is the functional equivalent of the centrosome in higher eukaryotic cells and the essential yeast spindle pole components have homologs in humans [1–5]. However, the yeast SPB exhibits a morphologically distinct structure from higher eukaryotic centrosomes. In contrast to the fluid matrix of the pericentriolar material built around the centrioles of mammalian centrosomes [6,7], the SPB exists as three highly organized, stratified layers built around a crystalline core of Spc42 [8–11]. Our understanding of SPB structure and composition has been advantageously used to understand highly conserved mechanisms of centrosomal regulation through phosphorylation [12–14]. Centrosomes serve as signaling platforms, integrating cell signals to regulate the localization of spindle proteins and progression through the cell cycle [5,15–22]. More specifically, kinases Mps1, Polo, Hrr25, and Cdk1 have been implicated in the regulation of SPB duplication, SPB separation, and cell cycle transitions [5,12,13,23–26].

Previous research shows that all SPB proteins are phosphoproteins. Two of the best characterized phosphoproteins at the SPB are Tub4 and Spc110, both implicated in microtubule nucleation. Several individual phosphorylation sites have been identified and mutated in Tub4. Phosphomimetic mutations of highly conserved Tub4 sites induce a mitotic arrest and confer defects in spindle assembly (S360) [27,28], increase microtubule assembly rates and numbers of microtubules at the SPB (Y445) [29], and induce metaphase arrests with short, disorganized spindles (S74 and S100) [28]. Likewise, Spc110 phosphorylation has been studied extensively *in vivo* [30–33]. Mps1 phosphorylation of Spc110 S60, T64, and T68 is responsible for a gel shift during mitosis [30–32] and blocking phosphorylation of S91 induces a metaphase delay [33].

Two large-scale analyses of SPB phosphorylation have been performed. Valuable identification of phosphorylation sites in the γ-tubulin small complex (γ-TuSC) relied on material overexpressed in yeast, which could alter the phosphorylation pattern [28]. Phosphoanalysis of purified SPBs identified 298 sites but sequence coverage of the γ-TuSC was poor compared to the coverage of the other components [27].

Using our improved SPB purification protocol [34] and relying on advances in mass spectrometers and analysis packages, we have conducted a new phosphoanalysis of purified SPBs. Our data expand on previously reported phosphoproteomic data sets and identify novel cell cycle phosphorylation states of the SPB at the onset of microtubule nucleation in G1/S, a cell cycle stage not previously examined. Our phosphorylation data of the microtubule nucleating γ-TuSC builds on the current literature by identifying and fluorescently analyzing new cell cycle phosphorylation sites of the specific population of γ-TuSCs attached to SPBs.

## RESULTS AND DISCUSSION

### Purification by TAP-tagged Spc97 increases the yield of intact purified spindle pole bodies

Previous studies have shown that SPBs from *Saccharomyces cerevisiae* co-purify with Mlp2, a nuclear pore component [35]. To increase the yield of purified SPBs, tandem affinity purification (TAP) tags were introduced on spindle pole components (List of strains in Table 1). Starting with the proteins found in the core of the SPB, we tagged Spc42 and Cnm67. While N-terminally tagged Cnm67 was viable, we found that the core spindle pole component Spc42 tagged with a C-terminal TAP tag was not viable in our strain background. In addition to core proteins, we tagged membrane anchor proteins Nbp1 and Bbp1, linker proteins Spc110 and Spc72, and γ-TuSC component Spc97 with C-terminal TAP tags.

**Table 1.**
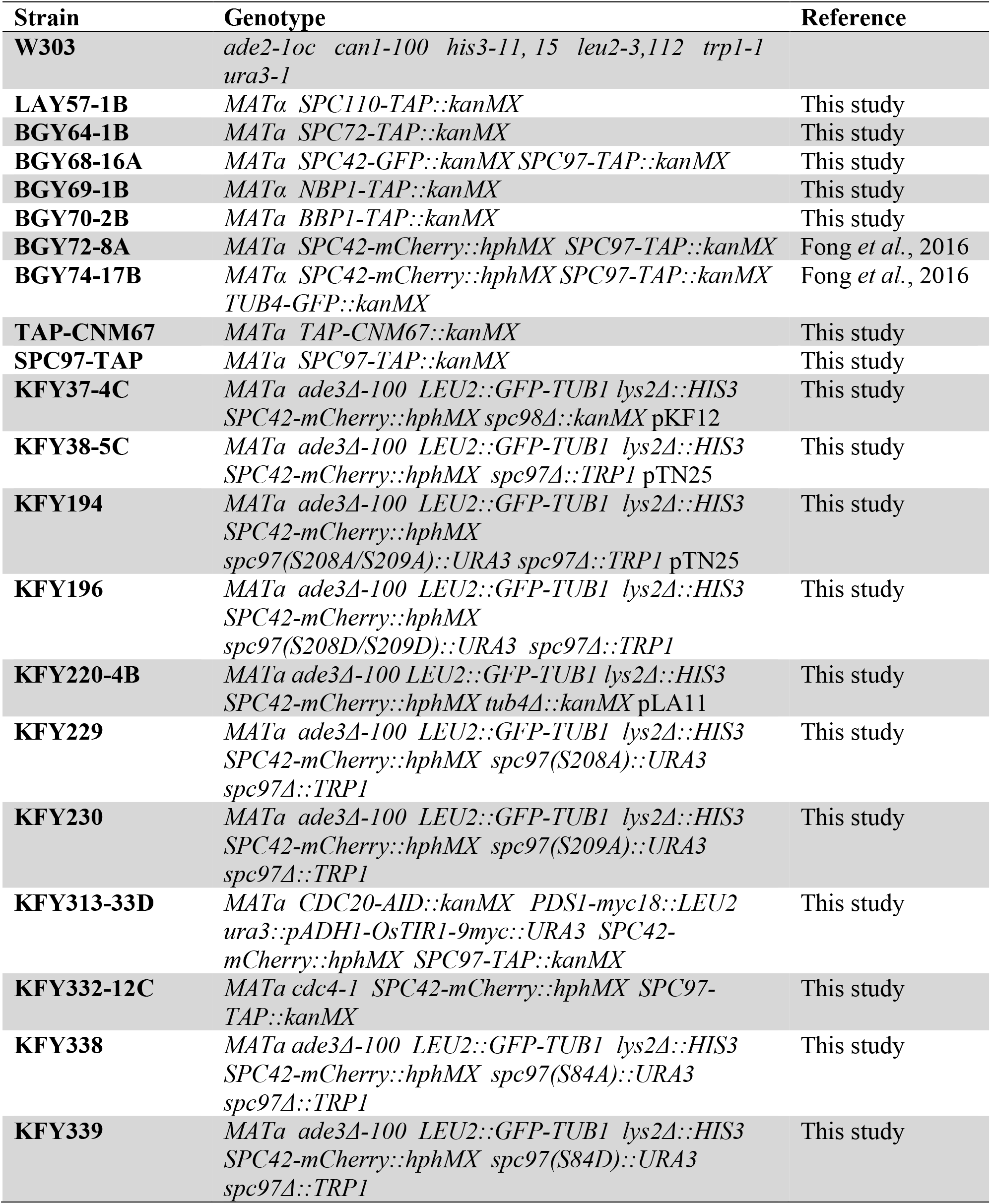
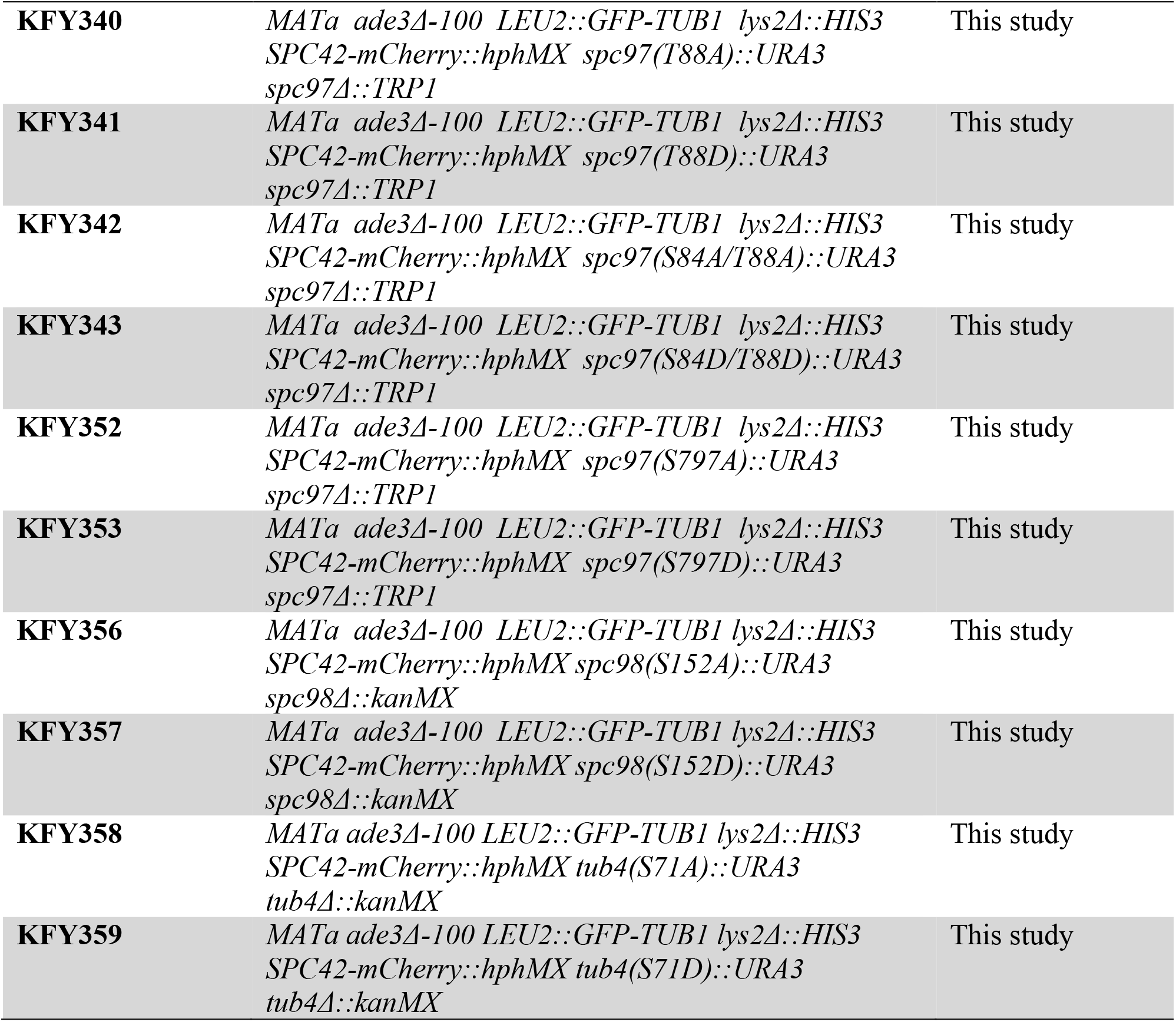
Strains used in this study.

We used western blot analysis to assess the yield of purified SPBs and found that Mlp2-TAP bound weakly to IgG beads, resulting in a low yield of purified SPBs. Similarly, Bbp1-TAP, Nbp1-TAP and Spc72-TAP showed weak or no binding to IgG beads. Other constructs (TAP-Cnm67 and Spc110-TAP) showed relatively strong binding to IgG beads, but TEV (tobacco etch virus) protease cleavage failed to remove the purified SPBs from the beads, again resulting in low yields. A C-terminal TAP tag on Spc97 was shown not only to bind the strongest to IgG beads, but to be efficiently cleaved from the beads by TEV protease, resulting in the highest, most reproducible yield of purified SPBs (Figure 1A).

**Figure 1.**
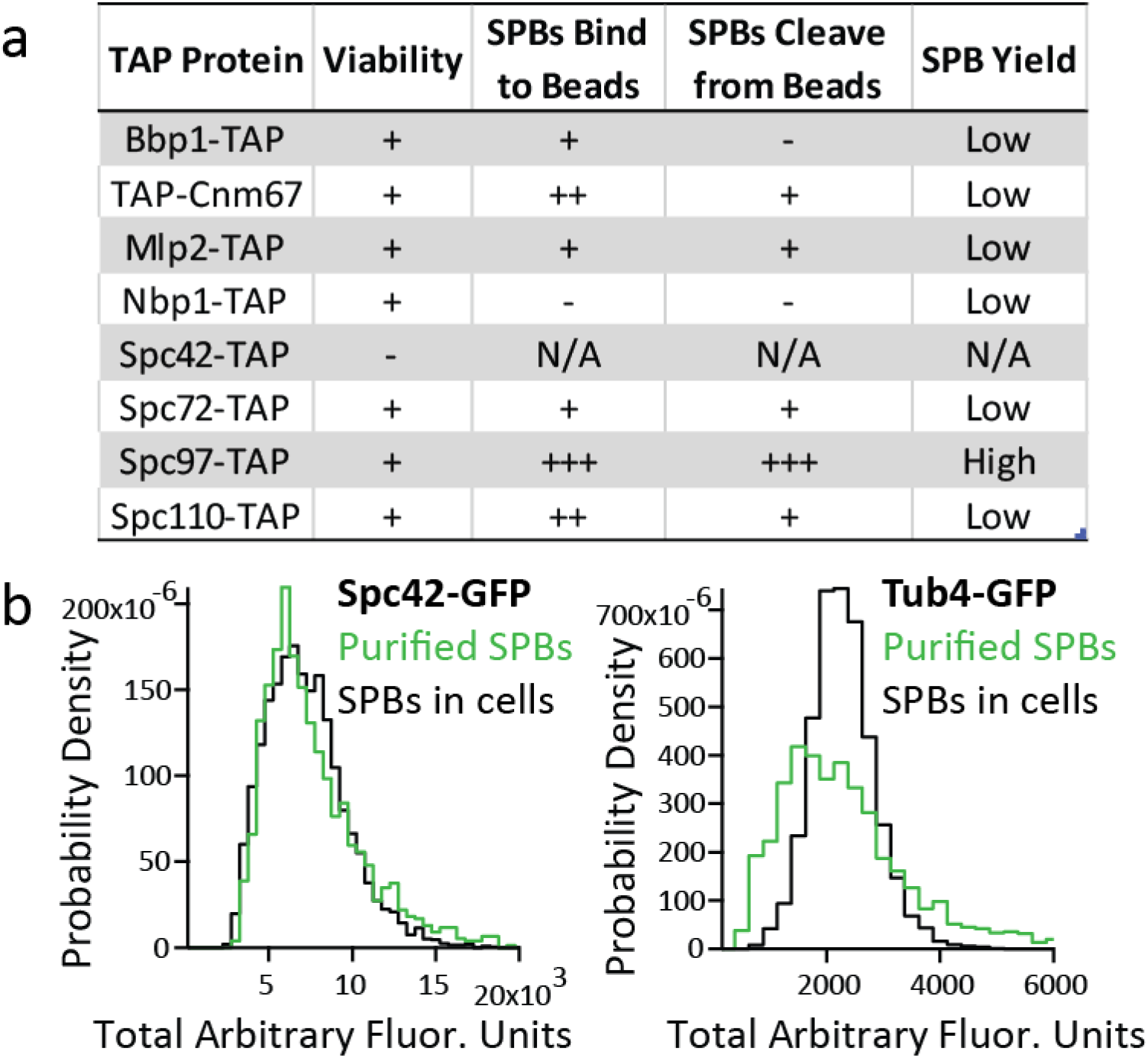
Purification protocol yields intact spindle pole bodies. a. Several components of the spindle pole body (SPB) were tagged with a TAP tag. SPBs were purified using each tag and the yield of SPBs determined. The efficiency of SPBs binding to beads and being cleaved from beads were determined by western blot analysis. b. Purified SPBs are the same size as SPBs in cells, as determined by fluorescence intensities of SPBs containing either Spc42-GFP or Tub4-GFP imaged in cells (in black) and after purification (in green).

Spc97 is a component of the γ-TuSC, a stable heterotetrameric complex that exists free in solution as well as bound to the SPB. Because our purification relied on a TAP tag on Spc97, our protocol yielded free γ-TuSCs in addition to intact SPBs. Separation of intact SPBs and γ-TuSCs was achieved by velocity sedimentation [34]. Analysis by western blot confirmed that SPBs were concentrated in fractions 9-11 of the sucrose gradient (40-50% sucrose) while the soluble fraction of the γ-TuSCs remained higher in the gradient [34].

### Purified spindle pole bodies are the same size as spindle pole bodies *in vivo*

To determine if purified SPBs were intact and the same size as *in vivo*, SPBs were purified from strains that contained either Spc42-GFP or Tub4-GFP. Quantification of the fluorescence intensities verified purified SPBs containing Spc42-GFP had the same fluorescence intensity as Spc42-GFP SPBs in live cells, suggesting that the core of the SPB remained intact through the purification. To test if the inner and outer plaques were retained during the purification, SPBs containing Tub4-GFP were purified and imaged *in vitro* and *in vivo*. Quantification of fluorescence intensity indicated that purified SPBs retained 88% of the γ-TuSC compared to SPBS in live cells (Figure 1B). Finally, all eighteen proteins were present as determined by high protein coverage in mass spectrometry (Table 3).

**Table 3.** Sequence coverage (%) of SPB proteins by mass spectrometry. Each column corresponds to a biological replicate. For each biological replicate, with the exception of the first asynchronous SPB sample, two technical replicates were individually analyzed by Comet [50], then combined for Percolator [51] and protein inference analyses. Three biological replicates were completed for asynchronous SPBs, three for mitotic SPBs, and two for G1/S SPBs. The percentage sequence coverage for each SPB protein is shown.

|  | Asynchronous SPBs |  |  | Mitotic SPBs<br>Cdc20-AID |  |  | G1/S SPBs<br>cdc4-1 |  |
| --- | --- | --- | --- | --- | --- | --- | --- | --- |
| **Bbp1** | 67 | 52 | 0 | 41 | 18 | 64 | 78 | 70 |
| **Cdc31** | 71 | 52 | 44 | 52 | 47 | 68 | 72 | 74 |
| **Cmd1** | 63 | 50 | 33 | 50 | 47 | 88 | 63 | 63 |
| **Cnm67** | 71 | 57 | 46 | 55 | 71 | 85 | 74 | 69 |
| **Kar1** | 63 | 40 | 0 | 29 | 29 | 47 | 69 | 61 |
| **Mps2** | 49 | 39 | 0 | 31 | 7 | 37 | 69 | 59 |
| **Mps3** | 49 | 29 | 2 | 24 | 20 | 60 | 59 | 51 |
| **Nbp1** | 53 | 40 | 0 | 32 | 21 | 50 | 53 | 52 |
| **Ndc1** | 35 | 20 | 0 | 17 | 19 | 39 | 55 | 38 |
| **Nud1** | 49 | 36 | 28 | 28 | 62 | 59 | 75 | 57 |
| **Sfi1** | 41 | 23 | 0 | 15 | 7 | 24 | 40 | 31 |
| **Spc29** | 76 | 49 | 14 | 60 | 69 | 71 | 76 | 68 |
| **Spc42** | 77 | 52 | 27 | 56 | 69 | 80 | 82 | 79 |
| **Spc72** | 58 | 40 | 42 | 32 | 67 | 69 | 69 | 61 |
| **Spc97** | 52 | 41 | 45 | 35 | 73 | 65 | 72 | 62 |
| **Spc98** | 42 | 35 | 29 | 29 | 52 | 58 | 61 | 49 |
| **Spc110** | 69 | 58 | 31 | 54 | 57 | 69 | 75 | 72 |
| **Tub4** | 46 | 10 | 20 | 21 | 62 | 61 | 68 | 52 |

### Tandem mass spectrometry data identified phosphorylation sites of the spindle pole body at different cell cycle stages

SPBs were purified from different stages of the cell cycle to identify cell cycle specific phosphorylation events. The phosphorylation state of SPBs at G1/S had not been previously described. We purified SPBs from *cdc4-1* cells, which were arrested in G1/S by a shift to the restrictive temperature of 36°C. To identify the subset of phosphorylation sites present during mitosis, SPBs were harvested from cells arrested in mitosis by depletion of Cdc20. Finally, we purified SPBs from asynchronous cultures. Western blot analysis after velocity sedimentation verified the presence of SPBs in fractions 9–11 of the sucrose gradient (40–50% sucrose) using antibodies against Spc110 and Spc97 (Figure 2A).

**Figure 2.**
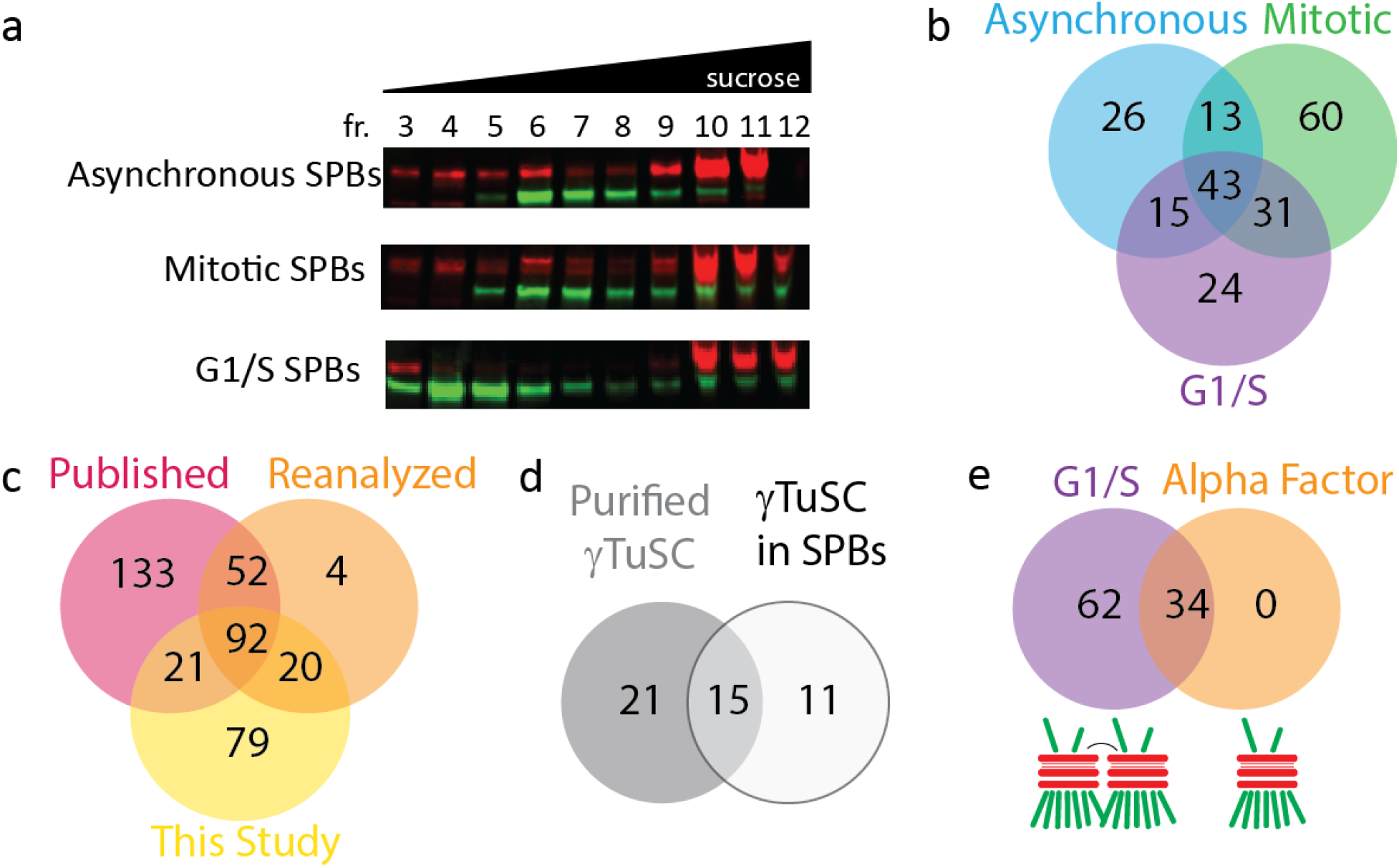
Comparison of phosphoproteomic data sets for the spindle pole body isolated from different cell cycle stages. a. Asynchronous, mitotic, and G1/S SPBs were purified. Fractions containing intact SPBs were determined by western blot analysis, using antibodies against Spc110 (in red) and Spc97 (in green). Sucrose concentration increases from left to right. Fractions with high concentrations of spindle pole bodies (Fractions 9, 10, 11 for asynchronous SPBs; Fractions 10, 11, for mitotic SPBs and G1/S SPBs) were used for phosphoproteomic mass spectrometry analysis. b. Comparison of high resolution phosphoproteomic data collected in this study for asynchronous wild type (in blue), mitotic (in green), and G1/S (in purple) SPBs. c. Comparison of the phosphoproteome published in Keck *et al.*, 2011 (in red), reanalysis of Keck *et al.*, 2011 data (in orange), and data collected in this study (in yellow). d. Comparison of γ-tubulin small complex phosphorylation sites identified in overexpressed γ-tubulin subcomplexes (Lin *et al.*, 2011; in grey) and identified in the context of the intact SPB (this study; in white). e. Comparison of G1 SPBs arrested by *cdc4-1* or alpha factor. In *cdc4-1* arrest, as indicated by the cartoon, the SPBs are duplicated, but not separated and both the new and the old SPBs are nucleating microtubules. In an alpha factor arrest, the SPB has not yet duplicated.

Phosphorylation sites were identified by mass spectrometry and verified manually to eliminate any ambiguous or unsubstantiated assignments. For the phosphoproteomic data set reported here, 212 phosphorylation sites were reported across asynchronous, mitotic, and G1/S SPBs (Figure 2B, Supplementary File). Of the sites reported in this study, 60 (28%) were only identified in mitotic SPBs, and 24 (11%) were only identified in G1/S SPBs and 26 (12%) were only detected in the asynchronous SPB sample. Forty-three sites (20%) were found in SPBs from all three conditions.

A previous study of the yeast SPB phosphoproteome reported 298 phosphorylation sites in the eighteen SPB components throughout the cell cycle [27]. However, since publication, advances in the resolution and sensitivity of mass spectrometers and new software packages that apply stringent quality cutoffs increased the confidence in phosphorylation assignments. Of the phosphorylation sites identified in this current study, 113 phosphorylation sites were also identified in Keck *et al*., 2011, resulting in 28% (113/397 total sites) of all detected sites appearing in both data sets. To determine if the difference between the data sets was due to how the mass spectra were analyzed, we reanalyzed the raw data from Keck *et al*., 2011 using Comet and Percolator, the same algorithms used for our new data set. The reanalysis of the Keck *et al*, 2011 raw data set resulted in a reduction from 298 published phosphorylation sites to 168 phosphorylation sites (for more about statistical analysis, phosphorylation assignments, reanalysis of Keck *et al*, 2011 data, and dataset comparisons, see Supplemental Table 1 and Supplemental Figures 1–12). Many ambiguous assignments were removed from the previously published data set by imposing strict statistical cutoffs for spectra included in the data set. The reanalysis of previously published data and our new data set showed 42% (112/268 total sites) agreement (Figure 2C, Supplemental File). The 56 sites identified in the reanalyzed published data, but not this data set, most likely result from the fact that the Keck *et al*, 2011 sample was enriched for phosphopeptides by running the sample over a titanium dioxide column [27]. In contrast, phosphopeptides were not enriched for this data set, thus some rare phosphorylation events were likely missed. The 100 sites only identified in this study might result from improved SPB purification or advances in mass spectrometer instrumentation and sensitivity.

Previous work focused on phosphorylation of the γ-TuSC overexpressed and purified from yeast [28]. Looking at the γ-TuSC phosphorylation sites in Spc97, Spc98, and Tub4 identified in our data set, we find that 15 of the sites agreed with the reported literature. However, our data has identified 11 additional sites, 6 of which were identified in G1/S SPBs, on γ-TuSC that might inform our understanding of the phosphorylation state specifically when attached to the SPB (Figure 2D).

### The phosphorylation profile of G1/S spindle pole bodies include sites present at the initiation of microtubule nucleation

Previous studies on SPB phosphorylation used alpha factor to arrest the cells in G1 [27]. We report a novel phosphorylation state of SPBs from cells arrested in G1/S with a *cdc4-1* temperature sensitive arrest. These two G1 arrests differ in the state of the SPB and in the active cyclins present. In an alpha factor arrest, the SPBs are not duplicated and Clb-Cdc28 is inhibited [36–38]. In contrast, in a *cdc4-1* temperature sensitive arrest, the SPBs are duplicated but not separated [39,40] and there are high levels of Cln-Cdc28 and low activity of Clb-Cdc28 [41,42]. The *cdc4-1* shift to the restrictive temperature results in the earliest cell cycle arrest in which the newly duplicated SPB has nucleated microtubules, suggesting that the phosphorylation profile at this arrest is the earliest state conducive to high microtubule nucleation activity. Interestingly, all phosphorylation events detected during an alpha factor arrest were also identified in a G1/S arrest. However, there were several additional G1/S phosphorylation events that were detected in SPBs at this stage of the cell cycle (Figure 2E).

In alpha factor arrested SPBs from Keck *et al*, 2011, very few phosphorylation sites were identified on the γ-TuSC (Spc97, Spc98 and Tub4) or the proteins that bind the γ-TuSC to the core of the spindle pole body (Spc72 and Spc110) [27]. In fact, reanalysis of the published data failed to identify any unique G1 phosphorylation events for any of these five proteins. In contrast, in this study, each of the three γ-TuSC proteins—Spc97, Spc98, and Tub4—contained phosphorylation sites only observed in G1/S SPBs as well as five G1/S phosphorylation sites on Spc110 (Supplemental File).

### Phosphorylation of the γ-tubulin small complex is required for establishment of a proper mitotic spindle length

We examined the role of γ-TuSC phosphorylation events on spindle morphology, focusing on sites predicted to interfere with interaction with Spc110. Spc97 S84 was identified in mitotic and G1/S SPBs and T88 was identified as a phosphorylation event in G1/S. Both of these sites were identified in free γ-TuSC, but not in previous phospho-analyses of intact SPBs [27,28]. These two sites map to the outer face of γ-TuSC, in a region predicted to interact with Spc110 (Figure 3A). Phosphomimetic and phosphoblocking mutations of these sites were integrated into fluorescent yeast strains (List of strains in Table 1; List of plasmids in Table 2).

**Figure 3.**
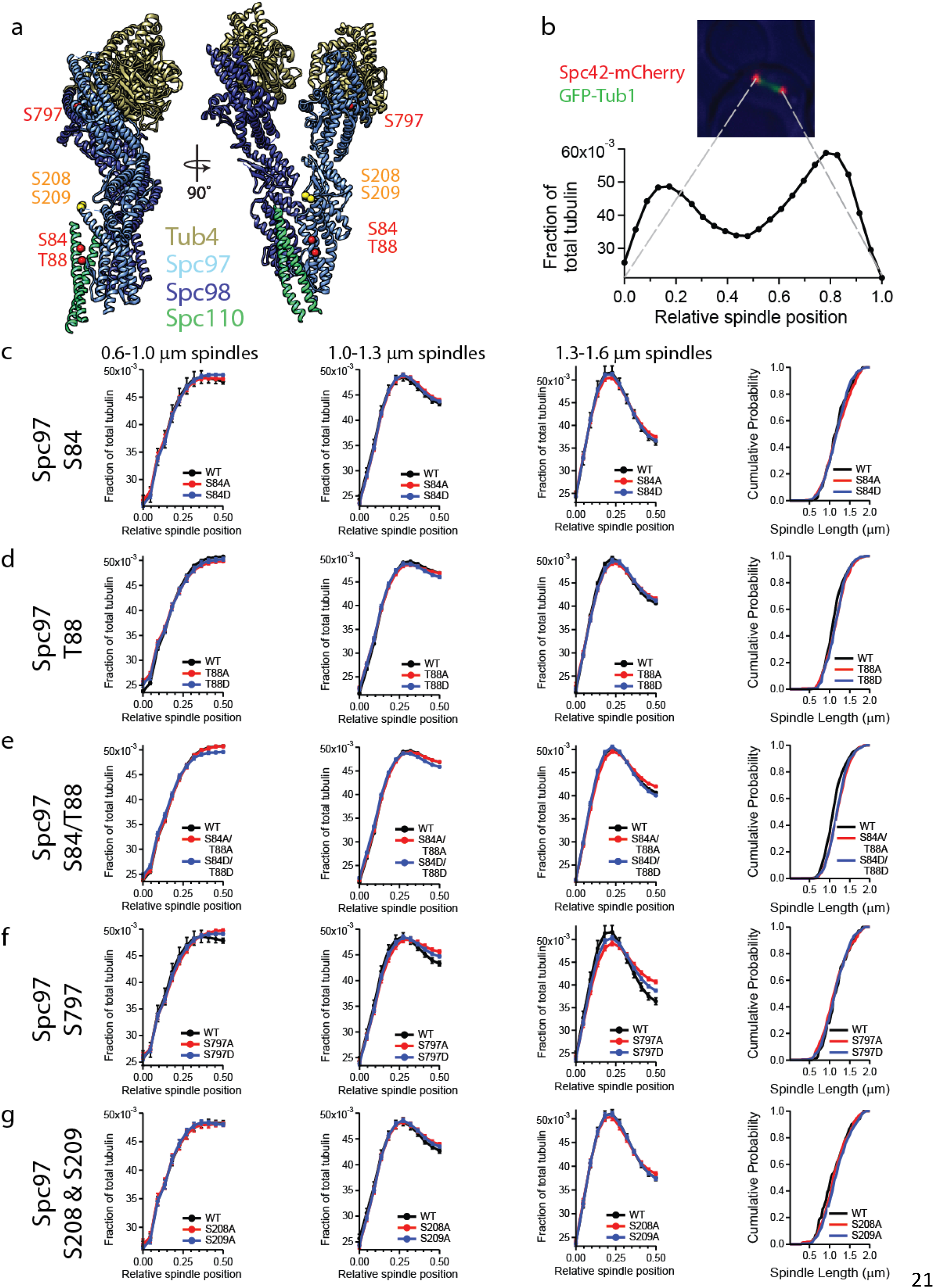
Phosphorylation of Spc97 is required for normal mitotic spindle morphology. a. Mitotic and G1/S phosphorylation sites (in red) have been mapped onto the pseudo-atomic structure of Spc97 (light blue) in the yeast γ-tubulin small complex, adapted from PDB:5FM1 [53]. Phosphoblocking mutations of Spc97 S208 and S209 (in yellow) in combination are lethal. b. Spindle morphology was determined by measuring spindle length and tubulin fluorescence distribution along the spindle. Metaphase spindles were identified as spindles with 1.3–1.6 μm between Spc42-mCherry spindle poles. GFP-Tub1 fluorescence was measured across the entire spindle. c-g. Half-spindle tubulin distribution profiles for spindles of a given length are shown for several Spc97 phosphorylation mutants. Spindle tubulin distributions were separated into two half spindles for ease of display. In each of the tubulin distribution graphs, the SPB is located at 0 and the spindle equator is located at 0.5. The far right panels show a cumulative probability of spindle length for the observed spindles, measured as the distance between Spc42-mCherry foci.

**Table 2.** Plasmids used in this study.

| **Plasmid** | **Relevant Markers** | **Reference** |
| --- | --- | --- |
| **pRS316** | *CEN6 ARSH4 URA3 Amp<sup>r</sup> f1 origin* | Sikorski and Hieter, 1989 |
| **pRS306** | *URA3 f1 origin* | Sikorski and Hieter, 1989 |
| **pLA11** | *TUB4 ADE3 LYS2 2μm origin* | This study |
| **pKF12** | *SPC98 ADE3 LYS2 2μm origin* | This study |
| **pKF56** | *SPC97 in pRS316* | This study |
| **pKF137** | *spc97(S208A/S209A) in pRS306* | This study |
| **pKF161** | *spc97(S208D/S209D) in pRS316* | This study |
| **pKF191** | *spc97(S208A) in pRS306* | This study |
| **pKF192** | *spc97(S209A) in pRS306* | This study |
| **pKF244** | *spc97(S84A) in pRS306* | This study |
| **pKF245** | *spc97(S84D) in pRS306* | This study |
| **pKF246** | *spc97(T88A) in pRS306* | This study |
| **pKF247** | *spc97(T88D) in pRS306* | This study |
| **pKF248** | *spc97(S84A/T88A) in pRS306* | This study |
| **pKF249** | *spc97(S84D/T88D) in pRS306* | This study |
| **pKF258** | *spc97(S797A) in pRS306* | This study |
| **pKF259** | *spc97(S797D) in pRS306* | This study |
| **pKF262** | *spc98(S152A) in pRS306* | This study |
| **pKF263** | *spc98(S152D) in pRS306* | This study |
| **pKF264** | *tub4(S71A) in pRS306* | This study |
| **pKF265** | *tub4(S71D) in pRS306* | This study |
| **pSB2065** | *3V5-IAA7-kanMX Amp<sup>r</sup> f1 origin* | gift from Sue Biggins* |
| **pTN25** | *SPC97 ADE3 2μm origin* | This study |
| **PaTEVCa** | *N-terminal TAP tag* | This study |

\* Fred Hutch Cancer Resource Center, Seattle, WA

These mutations supported the growth of cells and conferred little change in the organization of the spindle microtubules or the length of the spindles (Figure 3C-E). However, the cells did exhibit a cell cycle delay, presenting as an increased percentage of large budded cells (18-20%) as compared to wild-type cells, where 4.7% were large budded.

Similar analysis was conducted for Spc97 S797. Spc97 S797 was a novel phosphorylation site identified in mitotic SPBs and maps to the C-terminal outer face of the γ-TuSC, which could also interact with Spc110 (Figure 3A). While phosphomimicking and phosphoblocking mutations had no effect on spindle length, tubulin distribution was affected by these mutations, with an increase of tubulin at the spindle midzone, suggestive of misregulation of kinetochore microtubule length (Figure 3F).

Fluorescent spindle analysis was also performed for a G1/S site detected in Spc98 (S152) and a G1/S site detected in Tub4 (S71) (List of strains in Table 1; List of plasmids in Table 2). Spc98 S152 is a possible Cdk1 phosphorylation site, which was also observed in free γ-TuSC [28]. While fluorescence analysis revealed no change in tubulin distribution across spindles of any length, the phosphomimetic mutation S152D showed a cell cycle delay with an increase in the length of arrested spindles and an increase in large budded cells from 6.7% to 17.3% suggesting a checkpoint delay (Figure 4C). The novel G1/S phosphorylation site in Tub4 S71 sits at the interface between γ-tubulin and α-tubulin (Figure 4A). The phosphoblocking mutation S71A resulted in longer spindles at the arrest and a cell cycle delay, with 18.3% large budded cells compared to 5% in wild type cells (Figure 4D). Phosphomimicking mutation of S71 also showed a cell cycle delay, with 27.7% large budded cells compared to 5% in wild type cells; however, there was no change in spindle length distributions.

**Figure 4.**
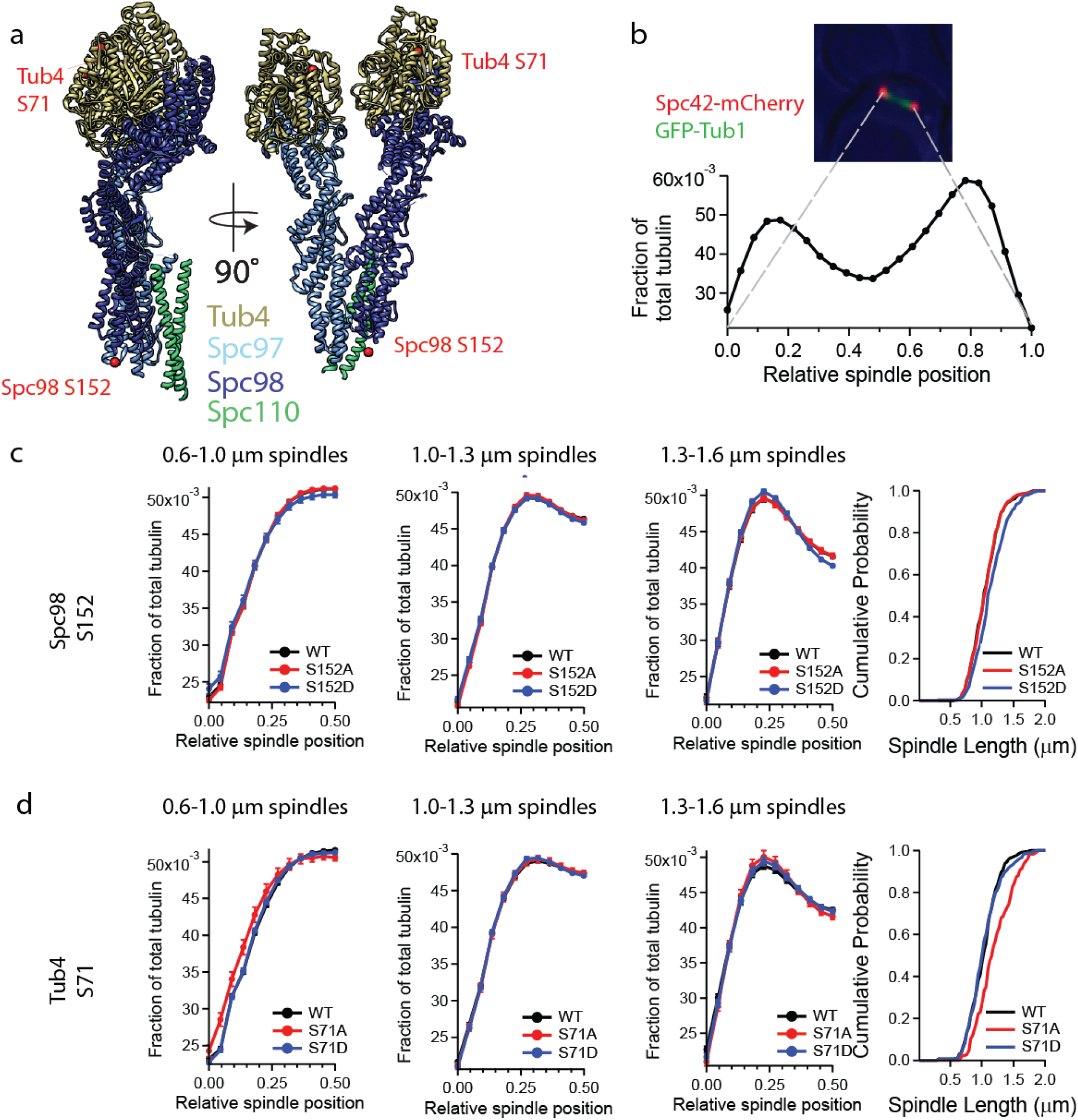
Phosphorylation regulation of the γ-tubulin small complex is required for normal mitotic spindle morphology. a. Phosphorylation site S71 is shown on the pseudo-atomic structure of Tub4 (gold) in the yeast γ-tubulin small complex, adapted from PDB:5FM1 [53]. The N-terminal 160 residues of Spc98 are not included in the pseudo-atomic structure, so the exact position of S152 is not shown. b. Spindle morphology was determined by measuring spindle length and tubulin fluorescence distribution along the spindle. Metaphase spindles were identified as spindles with 1.3–1.6 μm between Spc42-mCherry spindle poles. GFP-Tub1 fluorescence was measured across the entire spindle. c and d. Half-spindle tubulin distribution profiles for spindles of a given length are shown for Spc98 and Tub4 phosphorylation mutants. Spindle tubulin distributions were separated into two half spindles for ease of display. In each of the tubulin distribution graphs, the SPB is located at 0 and the spindle equator is located at 0.5. The far right panels show a cumulative probability of spindle length for the observed spindles, measured as the distance between Spc42-mCherry foci.

We also examined sites identified in mitotic SPBs, hypothesizing that these sites could be involved in regulation of microtubules during the establishment of the mitotic spindle. In previously published phosphoproteomic data, γ-TuSC component Spc97 was found to have two phosphorylation sites unique to mitosis, S208 and S209 [27]. Phosphomimetic mutations and phosphoblocking mutations were made for both mitotic phosphorylation sites and transformed into yeast (List of strains in Table 1; List of plasmids in Table 2). Using a plasmid shuffle assay, we determined that phosphomimicking mutations in Spc97 (S208D/S209D) were viable. However, the phosphoblocking mutations in Spc97 (S208A/S209A) were lethal. Spc97 S208A or Spc97 S209A were viable individually, with normal growth rates, morphologically wild type spindles and wild type tubulin distributions across metaphase spindles, suggesting that phosphorylation of at least one of these sites is required for progression through the cell cycle (Figure 3G). The locations of the Spc97 mitotic phosphorylation sites suggest that this region might be involved in interactions with Spc110. Disruption of the interface between Spc97 and Spc110 might destabilize the interaction between the γ-tubulin ring and Spc110.

## CONCLUSIONS

In summary, we optimized purification protocols to yield intact SPBs in quantities that facilitated *in vitro* analysis of SPB phosphorylation states by high-resolution mass spectrometry. While expanding on previously reported phosphorylation data sets, we were specifically interested in identifying phosphorylation sites in the γ-TuSC present at the SPB during G1/S arrest. We posit that that these G1/S sites reveal phospho-regulatory events important for the initiation of microtubule nucleation at the newly duplicated SPB. *In vivo*, mutation of G1/S and mitotic phosphorylation sites in the γ-TuSC illustrated the sensitivity of spindle organization and length regulation to perturbations at microtubule nucleation sites. This phosphoproteome provides a data set that will guide future work studying the regulation of the SPB and provide insight into the study of individual SPB components.

## MATERIALS AND METHODS

### Strains, plasmids, and media

The yeast strains used in this study were all derived from W303 and are listed in Table 1. C-terminally TAP tagged proteins were created by PCR amplifying the *TAP*-*kanMX* cassette from the plasmid TAP-2xPA using primers that shared homology with the flanking sequences of the stop codon in the gene of interest. Cnm67 was N-terminally TAP tagged by PCR amplification of the *TAP*-*kanMX* cassette from the plasmid PaTEVCa, using primers that shared homology with the flanking DNA of the start codon of *CNM67*. C-terminal mCherry and GFP protein fusions were created by PCR amplification of the *mCherry*-*hphMX3* and the *GFP*-*kanMX* cassettes from pBS35 and pFA6-GFP(S65T)::kanMX plasmids, respectively (gifts from the Yeast Resource Center, University of Washington, Seattle, WA). The cassettes for the fluorescent proteins shared homology with the flanking sequence before the stop codon of the genes of interest. For the Cdc20-AID strain, the auxin degron IAA7 was PCR amplified from pSB2065 (gift from Sue Biggins, Fred Hutchinson Cancer Research Center, Seattle, WA) with primers that shared homology with the flanking DNA of the stop codon of the gene. The above cassettes were integrated into a diploid strain, KGY315, and verified by PCR.

The plasmids used in this study are listed in Table 2. QuikChange Lightning Multi Site-Directed Mutagenesis (Stratagene) was used to construct plasmids containing point mutations. Plasmids carrying mutations in *SPC97* were integrated at the *SPC97* locus in strain KFY38-5C (*spc97*Δ), plasmids carrying mutations in *SPC98* were integrated at the *SPC98* locus in KFY37-4C (*spc98*Δ), and plasmids carrying mutations in *TUB4* were integrated at the *TUB4* locus in KFY220-1A (*tub4*Δ) using a plasmid shuffle[43]. These three strains all contained GFP-Tub1 for imaging of microtubules and Spc42-mCherry to visualize spindle pole bodies.

YPD media is as described [44]. SD-ura low ade and SD-lys were previously described [45,46].

### TAP purification and velocity sedimentation

Spindle pole bodies were purified using a TAP-tag on Spc97, as previously described [34]. Sucrose gradients were generated by allowing five steps of sucrose solutions (200 µl each of 10%, 20%, 30%, 40%, 2.5 M sucrose) to equilibrate at 4°C for 2 hours. For fluorescence analysis, the sucrose solutions were made in 10 mM Bis-Tris, pH 6.5, 0.1 mM MgCl_2._ For mass spectrometry analysis, the sucrose solutions were made in 40 mM HEPES, pH 7.4, 150 mM NaCl. The TEV eluate was then applied to the sucrose gradient and spun at 50000 rpm for 5 hours at 4°C in a TLS55 rotor (Beckman Coulter). SPBs isolated from *cdc4-1* cells were spun at 50000 rpm for 4 or 4.5 hours to prevent side-by-side SPBs from settling to the bottom of the sucrose gradient. Fractions (90 µl) were removed from the top of the gradient with wide-bore tips. The presence of SPBs was determined by western blot analysis, probing for Spc110 and Spc97. Western blot analysis showed the separation of intact SPBs (fractions 9-11) from the soluble pool of spindle pole body components (fractions 1-8).

For isolation of SPBs from cells arrested by depletion of Cdc20, a strain carrying the Cdc20-AID was grown to 80 Klett units in YPD. Auxin (indole-3-acetic acid; IAA) in DMSO was added to a final concentration of 1 mM. Cdc20 was depleted for 1.5 generations before cells were harvested. In Cdc20 depleted cells, ≥98% of cells arrested with large buds. For isolation of SPBs from cells arrested at G1/S, a strain carrying *cdc4-1* was grown at 25°C to 60 Klett units, then shifted to the restrictive temperature of 36°C for two generations before harvesting cells. 95% of cells had elongated buds and 5% had large buds as expected for a *cdc4-1* arrest. Spindle pole bodies were then purified as previously described [34].

### Fluorescence microscopy

All images were acquired using a DeltaVision system (Applied Precision) with an IX70 inverted microscope (Olympus), a U Plan Apo 100x objective (1.35 NA) and a CoolSnap HQ digital camera (Photometrics). Exposures were 0.4 s for mCherry and GFP. Images were processed as previously described [47] using custom Matlab programs (Fluorcal and Calcmate) to identify GFP and mCherry foci and quantify the fluorescence intensities. Fluorcal and Calcmate are available upon request. For live-cell imaging, cells were mounted on an agarose pad as previously described [48]. Metaphase spindles were identified as spindles with 20-25 pixels (1.3-1.6 μm) between spindle poles.

For *in vitro* SPB imaging, a flow cell was constructed with KOH cleaned glass coverslips. Spindle pole bodies were diluted with 5X BRB80/BSA (400 mM K-PIPES, pH 6.9, 5 mM MgCl_2_, 5 mM EGTA, 40 mg/ml BSA) and KCl to a final concentration of BRB80, 8 mg/ml BSA and 500 mM KCl. The diluted SPBs were flowed into the flow cell and allowed to nonspecifically adhere to the coverslip for 30 minutes before imaging.

### Mass spectrometry sample preparation and digestion

Purified SPB fractions in 40–50% sucrose, 40 mM HEPES, pH 7.4, and 150 mM NaCl, were combined such that samples were approximately 2 to 30 μg total protein in 0.5 to 1.5 mL of buffer plus approximately 45% sucrose. Samples were diluted 1:1 using 25 mM ammonium bicarbonate (ABC) and concentrated down to 30 μL using Amicon® Ultra 0.5 mL Centrifugal Filters with a 10,000 NMWL (Merc Millipore Ltd.) according to the manufacturer’s instructions. 500 μL of 25 mM ammonium bicarbonate was added to the top of the filter unit and spun through. This was repeated a total of three times. Sample volume was made up to 100 µL of 25 mM ABC and reduced in the filter unit with 10 mM dithiothreitol (DTT) at 37°C for 30 minutes followed by a 30-minute alkylation at room temperature with 15 mM iodoacetamide. 1 μL of 0.8 μg/μL Sequencing Grade Modified Trypsin (Promega Corporation) was used to digest the samples in the filter units at 37°C for 4 hours at 1,200 rpm in an Eppendorf Thermomixer. After digestion, peptides were spun through the filter units into a new Amicon Eppendorf tube. 100 μL of 25 mM ABC was added to the top of the filter unit and spun through into the same tube. The remaining digested sample was transferred from the filter unit into the Eppendorf tube by pipette. The digested sample was reduced to about 50 μL in a Speedvac. Sample pH was adjusted to 2 with 5 M HCl prior to storage at −80°C until mass spectrometry analysis.

### Mass spectrometry

Mass spectrometry was performed on either a Q-Exactive or Q-Exactive HF (Thermo Fisher Scientific). 3 μL of sample digest was loaded by autosampler onto a 150-μm Kasil fritted trap packed with Jupiter C12 90 Å material (Phenomenex) to a bed length of 2 cm at a flow rate of 2 μL/min. After loading and desalting using a total volume of 8 μL of 0.1% formic acid plus 2% acetonitrile, the trap was brought on-line with either a pulled fused-silica capillary tip (75-μm i.d.) or an empty Pico-Frit column (New Objective) that was self-packed with 30 cm of Reprosil-Pur C18-AQ (3-μm bead diameter, Dr. Maisch) mounted in an in-house constructed microspray source and placed in line with a Waters Nanoacquity binary UPLC pump plus autosampler. Peptides were eluted off the column using a gradient of 2-35% acetonitrile in 0.1% formic acid over 120 minutes, followed by 35-60% acetonitrile over 10 minutes at a flow rate of 250 nL/min.

The Q-Exactive mass spectrometer was operated using data dependent acquisition (DDA) where a maximum of twenty MS/MS spectra were acquired per MS spectrum (scan range of m/z 400 to 1600). The resolution for MS and MS/MS was 60,000 and 15,000, respectively, at m/z 200. The automatic gain control (AGC) targets for MS and MS/MS were set to 3e6 and 1e5, respectively, and the maximum fill times were 50 and 25 msec, respectively. The MS/MS spectra were acquired using an isolation width of 1.6 m/z and a normalized collision energy (NCE) of 27. The underfill ratio was set to 10% and the intensity threshold set to 4e5. MS/MS acquisitions were prevented for unassigned, +1, +6 and greater precursor charge states. Dynamic exclusion (including all isotope peaks) was set for 5 or 10 seconds. The Q-Exactive HF was operated similarly.

### Analysis of mass spectrometry data

Mass spectra from this study and from previously published data were converted into mzML using msconvert from ProteoWizard [49]. With the exception of one asynchronous sample, two technical replicates were run for each biological sample. For each technical replicate, proteins were identified by searching high-resolution MS/MS spectra against the SGD protein sequence database using Comet [50]. A variable modification of 79.966331 on S, T or Y was included in the search to identify phosphopeptides. Peptide identifications for the two technical replicates were then combined processed with Percolator [51]. MSDaPl was used to visually inspect the results [52] and confirm there was enough MS2 spectral evidence to differentiate between possible phosphorylated residues in a given peptide and give an accurate assignment. Because data dependent acquisition results in irreproducible sampling collection additional mass spectrometry runs might reveal more phosphorylation events at each stage of the cell cycle.

## ETHICS STATEMENT

No human or animal subjects were used in this study.

## DATA ACCESSIBILITY

Analysis of mass spectrometry data can be found at <u>http://www.yeastrc.org/fong_spb_2018</u>. The compiled datasets supporting this article have been uploaded as part of the supplementary material.

## COMPETING INTERESTS

The authors declare no competing interests.

## AUTHORS’ CONTRIBUTIONS

KKF and BG constructed strains and carried out protein purifications; KKF completed the fluorescence microscopy; JH constructed strains and completed fluorescence microscopy; AZ and RJ collected the mass spectrometry data; KKF, AZ, and MR participated in data analysis; KKF, MJM and TND drafted the manuscript and TND conceived and designed the study.

## ACKNOWLEDGEMENTS

We would like to thank the Yeast Resource Center for gift of plasmids (pBS35 and pFA6-GFP(S65T)::kanMX) for mCherry and GFP tagging, Sue Biggins (Fred Hutch Cancer Research Center) for gift of an auxin inducible degron tagging plasmid (pSB2065) and Gabrielle Grinslade for microscopy of Spc97 S84 and S797 mutants.

## FUNDING

This work was funded by NIH P01 GM105537 to Mark Winey and NIH P41 GM103533 to Michael MacCoss

